# Silent recognition of flagellins from human gut commensal bacteria by Toll-like receptor 5

**DOI:** 10.1101/2022.04.12.488020

**Authors:** Sara J. Clasen, Michael E. W. Bell, Du-Hwa Lee, Zachariah M. Henseler, Andrea Borbón, Jacobo de la Cuesta-Zuluaga, Katarzyna Parys, Jun Zou, Nicholas D. Youngblut, Andrew T. Gewirtz, Youssef Belkhadir, Ruth E. Ley

## Abstract

Flagellin, the protein unit of the bacterial flagellum, stimulates the innate immune receptor Toll-like receptor (TLR)5 following pattern recognition, or evades TLR5 through lack of recognition. This binary response fails to explain the weak agonism of flagellins from commensal bacteria, raising the question of how TLR5 response is tuned. Here, we describe a novel class of flagellin-TLR5 interaction, termed silent recognition. Silent flagellins are weak agonists despite high affinity binding to TLR5. This dynamic response is tuned by TLR5-flagellin interaction distal to the site of pattern recognition. Silent flagellins are produced primarily by the abundant gut bacteria *Lachnospiraceae* and are enriched in non-Western populations. These findings provide a mechanism for the innate immune system to tolerate commensal-derived flagellins.

**One-Sentence Summary:** TLR5 sensitively recognizes, but responds weakly to, flagellins from gut commensal bacteria.

## Main Text

Innate immune responses are initiated by pattern recognition receptors (PRRs) that evolved to detect conserved microbe-associated molecular patterns (MAMPs) (*1*). The Toll-like family of receptors (TLRs) are membrane-bound PRRs, widely expressed in many cell types, that activate pro-inflammatory pathways following MAMP-binding to their horseshoe-shaped ectodomains (*2*). Since MAMPs are not unique to pathogens, a question that has persisted for decades is whether TLRs respond differently to ligands derived from beneficial or commensal microbiota, relative to those produced by potentially pathogenic microbes (*3*). This question is especially relevant for TLRs that interface with the intestinal microbiota such as Toll-Like Receptor 5 (TLR5), which is highly expressed by epithelial cells that line mucosal surfaces (*4*).

TLR5 is plasma membrane-bound and binds extracellular flagellin, the protein subunit of the bacterial flagellum (*5*). Phylogenetically diverse bacteria produce structurally similar flagellins that consist of conserved N- and C-terminal D0-D1 domains separated by a hypervariable region (Fig. 1A) (*6*). The MAMP recognized by TLR5 is located in the N-terminal D1 (nD1) and referred to as the TLR5 epitope (*7, 8*). Studies on the FliC flagellin derived from the human pathogen *Salmonella enterica* serovar Typhimurium showed that mutating key residues in this region (FliC PIM) reduces ligand potency by several orders of magnitude (Table S1) and abolishes bacterial motility (*7*); crystal structures of FliC in complex with *Danio rerio* TLR5 later confirmed a direct interaction between these residues and the N-terminal region of the receptor ectodomain (*9*). Furthermore, flagellins that do not stimulate TLR5, like FlaA from the human pathogen *Helicobacter pylori* (‘*Hp*FlaA’), have different amino acids in their TLR5 epitope site (*8, 10*). TLR5’s inability to respond to *Hp*FlaA is characterized as ‘evasion’ and is presumed to occur through loss of TLR5 binding. Taken together, these studies demonstrate that robust TLR5 signaling requires the receptor ectodomain to bind the flagellin TLR5 epitope.

**Fig. 1.**
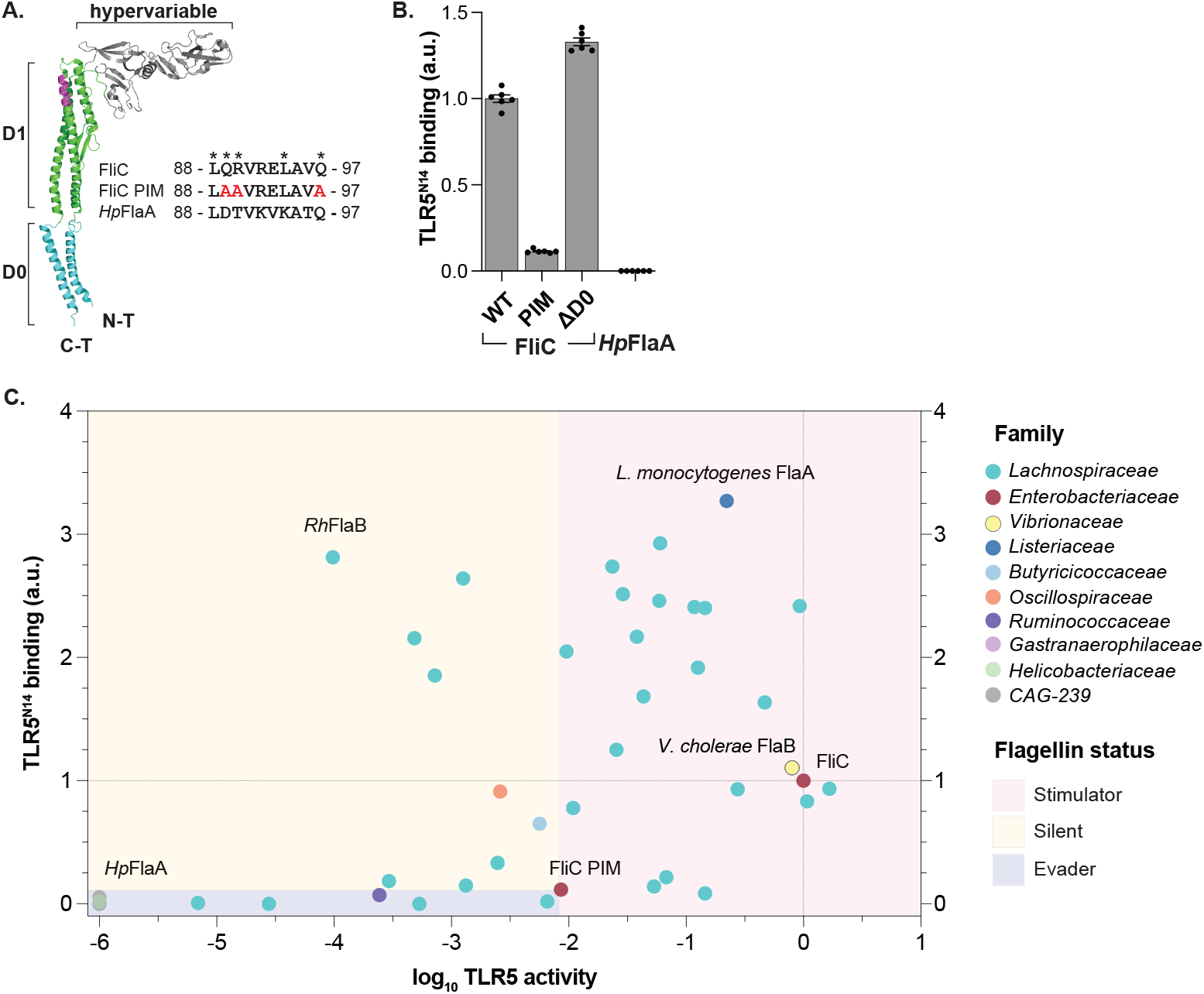
Flagellins from human gut commensals are silently recognized by TLR5. (**A**) Crystal structure of FliC (PDB 3A5X from (*6*)) and multiple sequence alignment of nD1 TLR5 epitope (colored magenta in structure) from *Salmonella* and *H. pylori* flagellins. Asterisks denote residues in FliC required for TLR5 recognition; residues mutated in FliC PIM are colored red. (**B**) Flagellin binding to truncated TLR5 ectodomain: TLR5^N14^ bait was incubated with AP-tagged flagellins followed by quantification of AP activity. Error bars are SEM for *n*=6; data shown represent two independent experiments. (**C**) Plot of TLR5 activity vs TLR5^N14^ binding for 41 flagellins abundant in the healthy human gut microbiome (see Fig. S1) as well as flagellins from pathogens. Circles represent individual flagellins and are colored by family-level taxonomy (GTDB). TLR5^N14^ binding was performed as described in B; data shown represent mean for *n*≥3. TLR5 activity was measured using TLR5 HEK-Blue cells and represents negative EC_50_ normalized to flagellin expression in bacterial lysates. Data represent mean from three independent experiments. All values are normalized to FliC. Flagellin status is defined relative to FliC PIM: stimulators are more active, silent flagellins are less active with higher affinity for TLR5^N14^, and evaders are less active with lower affinity for TLR5^N14^.

We quantified TLR5 recognition of flagellin by measuring the relative binding strength between the receptor and the nD1 epitope using a truncated form of the human ectodomain, TLR5^N14^ (similar to the one used in the crystal structure complex in (*9*)). This construct contains the first 14 leucine-rich repeats (LRRs) of the 22 LRRs that compose the ectodomain, including the flagellin nD1 binding site identified in the crystal structure, flanked by an N-terminal cap and C-terminal adaptor sequence tagged to IgG-Fc. Binding was quantified by incubating TLR5^N14^ with flagellins expressing C-terminal alkaline phosphatase (AP) and measuring AP activity. Consistent with its TLR5 epitope directly interacting with TLR5^N14^ (*9*), FliC binds strongly (Fig. 1B). In contrast, FliC PIM, which lacks three conserved residues in the epitope, shows a dramatic reduction in binding compared to FliC. Thus, the FliC-TLR5^N14^ interaction is primarily mediated by the nD1 TLR5 epitope, although a binding interface has been reported for the FliC cD1 domain (*9*). The D0 domain of FliC has previously been characterized as unnecessary for binding to TLR5 (*9, 11*). Consistent with the prediction that the D0 would not interact with TLR5, and with its retention of the nD1 epitope, we observed strong binding of FliC ΔD0 to TLR5^N14^. *Hp*FlaA fails to bind TLR5^N14^, congruent with its altered TLR5 epitope and reported lack of activation (*8, 12*) (Fig. 1A,B). These results indicate that TLR5^N14^ binding to flagellin reflects pattern recognition by TLR5 (*10*).

Flagellins have been characterized as either stimulatory (binding TLR5, leading to activation of the receptor, *e*.*g*., FliC), or evasive (no TLR5 activation, *e*.*g*., *Hp*FlaA) with the underlying assumption that TLR5 binding leads to activation of the receptor. However, flagellins from commensal bacteria induce a range of TLR5 activity (*13*–*15*), raising the question of how the TLR5 response to these flagellins is tuned. To investigate how TLR5 interacts with flagellins from commensal bacteria, we first searched for flagellins commonly encoded by the healthy human gut microbiome. Flagellin diversity is vast: of the 10 million proteins encoded by the human gut microbiome, over 5,000 different proteins are classified as flagellins (Methods). The majority of flagellin in the healthy human gut is produced by *Lachnospiraceae* (*16*), a prevalent and abundant family of *Firmicutes* that includes beneficial bacteria such as the butyrate-producers of the *Roseburia* and *Eubacterium* genera (*17*). Selecting from the most abundant flagellins observed in 270 healthy individuals (*18*) (Fig. S1), we expressed an initial 41 recombinant flagellins (34 belonging to *Lachnospiraceae* species) and screened these for both TLR5 signaling and TLR5^N14^ binding (Fig. S2A,B). Most of the 41 selected flagellins have TLR5 epitopes whose key residues are either identical to those of FliC (21/41) or differ at only one position (16/41) (Fig. S2C). In addition to flagellins from commensals, we included three flagellins from pathogens (FliC, *Vibrio cholerae* FlaB, and *Listeria monocytogenes* FlaA) and two negative controls (*Hp*FlaA and FliC PIM). We generated AP-tagged flagellins to assay TLR5^N14^ binding and separately expressed N-terminal Myc-tagged flagellins to quantify TLR5 activation. Flagellins were incubated with NF-kB reporter HEK cells engineered to express TLR5 and NF-kB-dependent AP activity was measured as a readout for TLR5 activation.

Consistent with the notion that binding TLR5 leads to its activation, we generally observed a positive relationship between TLR5^N14^ binding and TLR5 activity (Fig. 1C). Flagellins that induce a greater response than that of FliC PIM we categorized as ‘stimulators’ (red region in Fig 1C), regardless of their ability to bind TLR5^N14^; this describes nearly half the flagellins in our screen. ‘Evaders’, in contrast, bind and stimulate more weakly than FliC PIM (blue region). This group includes *Hp*FlaA and 12 commensal-derived flagellins. The remaining flagellins (9/41) resemble evaders with respect to TLR5 activation (stimulate worse than FliC PIM) but act like stimulators with regard to TLR5^N14^ binding (stronger than FliC PIM; yellow region). We termed these unexpected ligands ‘silent’ flagellins in reference to their inability to induce signaling despite intact TLR5 recognition.

We further investigated how silent flagellins decouple TLR5 ectodomain binding from agonism. We selected FlaB from *Roseburia hominis* (*Rh*FlaB) as our representative silent flagellin because it binds TLR5^N14^ the strongest among the silent flagellins from our initial screen (Fig. 1C). *R. hominis* is of wide interest, as it is a common gut commensal species belonging to the *Lachnospiraceae* and generally thought to be anti-inflammatory and thus beneficial to host health (*19*). Previous work demonstrated that *R. hominis* is motile and expresses *Rh*FlaB *in vivo (19, 20)*. We purified recombinant *Rh*FlaB and observed that, in addition to binding TLR5^N14^, it also binds full-length human TLR5 (Fig. 2A; lane 5). Of note, FliC binds more full-length TLR5 compared to *Rh*FlaB, while the evader *Hp*FlaA shows an equal lack of affinity for both truncated and full-length TLR5 (Fig. 2A). We also validated that *Rh*FlaB is a weaker TLR5 agonist than FliC PIM, despite its intact TLR5 epitope (Fig. 2B; Table S1).

**Fig. 2.**
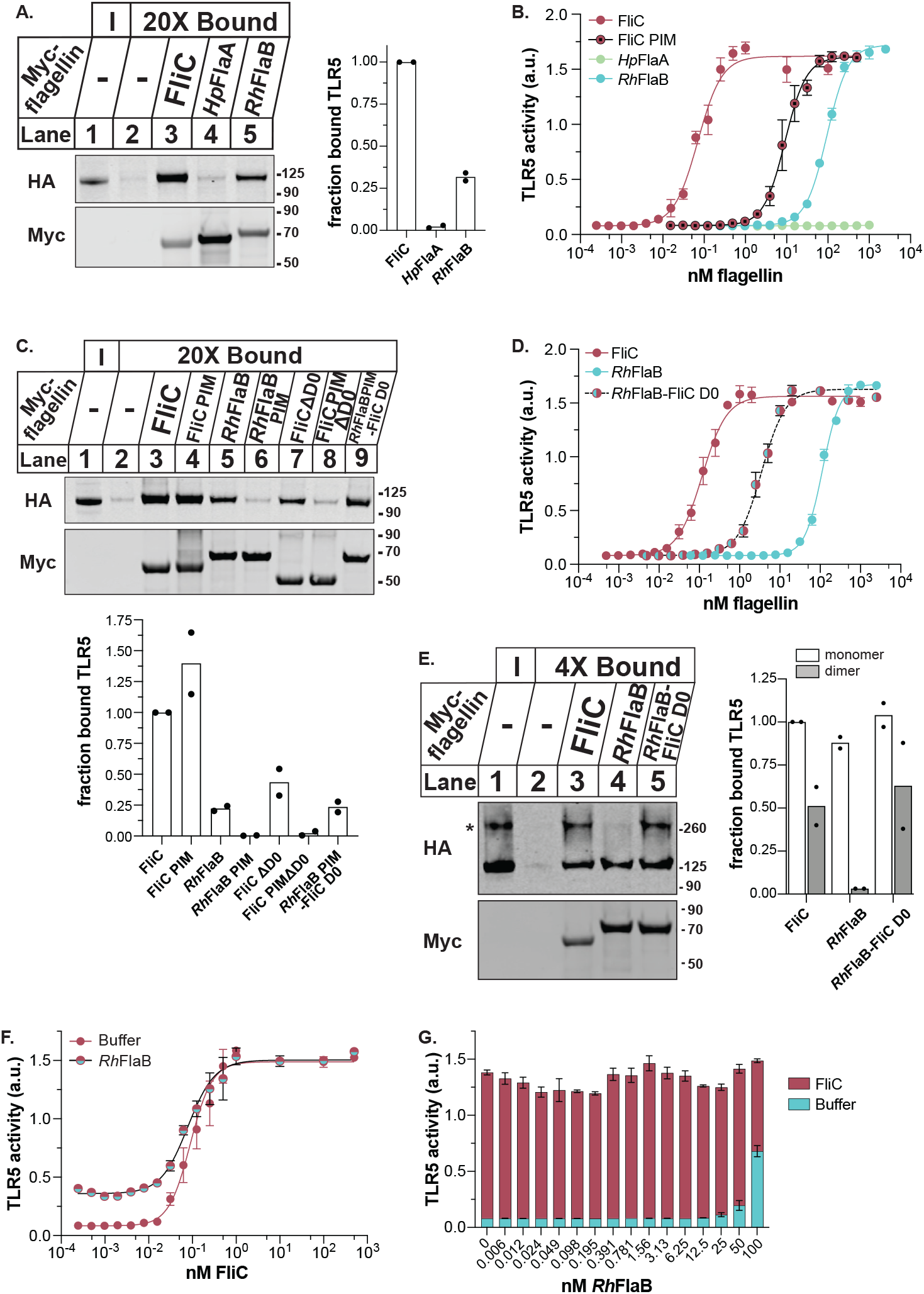
Silent flagellin *Rh*FlaB lacks TLR5 binding site in D0. (**A**) Flagellin binding to full-length TLR5: TLR5-HA HEK cell lysates were incubated with 6xHis-Myc-tagged flagellins followed by purification on Talon beads. Input (‘I’) and bound fractions (‘20X Bound’) were analyzed by immunoblot using antibodies against HA and Myc. *Left*: representative blots from one of two independent experiments; *right*: quantification of HA signal in bound lanes relative to Myc signal, normalized to FliC. (**B**) *Rh*FlaB-dependent TLR5 activity: TLR5 HEK-Blue cells were incubated with purified recombinant flagellins for 18 hr and NF-kB-dependent AP levels in medium were quantified. Error bars are SEM for *n*=3; data shown represent one of at least two independent experiments. Curve-fitting by weighted, non-linear regression analysis. (**C**) Mapping TLR5 binding sites in flagellin: TLR5-HA HEK cell lysates were incubated with 6xHis-Myc-tagged flagellins and processed as described in (A). *Top*: representative blots from one of two independent experiments; *bottom*: quantification of HA signal relative to Myc signal. (**D**) *Rh*FlaB chimera-dependent activation of TLR5: TLR5 HEK-Blue cells were incubated with purified recombinant flagellins as described in (B). (**E**) Flagellin binding to preformed TLR5 complexes: TLR5-HA HEK cells were treated with BS^3^ crosslinker prior to lysis then processed as described in (A). Asterisk indicates TLR5 dimer band. Quantification normalized to HA monomer bound to FliC. (**F, G**) FliC-dependent TLR5 activity in the presence of *Rh*FlaB: TLR5 HEK-Blue cells were exposed to 10 nM *Rh*FlaB (F) or 50 pM FliC (G) and buffer controls (superimposed in G) prior to the addition of flagellins at indicated concentrations and processed as described in (B).

Next, we tested if *Rh*FlaB binds TLR5 through its TLR5 epitope. We constructed the flagellin *Rh*FlaB PIM, which carries the same mutations as FliC PIM that result in loss of binding to TLR5^N14^. *Rh*FlaB PIM fails to bind the full-length receptor, consistent with TLR5 binding occurring solely at the TLR5 epitope (Fig. 2C; lanes 5-6). However, unlike *Rh*FlaB PIM, FliC PIM shows no reduction in binding to full-length TLR5 (Fig. 2C; lanes 3-4). This result was unexpected, because FliC PIM does not bind TLR5^N14^ ((*9*), and Fig. 1B).

We hypothesized that FliC PIM binds the C-terminal LRRs of the TLR5 ectodomain at a location allosteric to the site of pattern recognition. While the structure of this region of TLR5 remains unsolved, the C-terminal LRRs are predicted to interact with the conserved D0 domain of flagellin (Fig. 1A) (*21*). Notably, the D0 domain is not required for binding TLR5^N14^ (Fig. 1B) and is also absent in the FliC-TLR5 crystal structure (*9*). Several studies previously reported the necessity of the FliC D0 for TLR5 activation (*9, 11*). However, the mechanism is unclear and the authors unequivocally concluded that the D0 domain does not directly bind the receptor.

We tested for a TLR5 binding site in the FliC D0 using FliC PIM ΔD0 and assessing its binding to full-length TLR5. Since FliC PIM does not bind TLR5^N14^, if the D0 binds TLR5 LRRs 15-22, then FliC PIM ΔD0 should be unable to bind full-length TLR5. Consistent with an additional binding site in the D0 of FliC, FliC PIM ΔD0 shows a substantial loss of binding to TLR5 compared to FliC PIM and FliC ΔD0 (Fig. 2C; lane 8 vs lanes 4,7). Given our observation that *Rh*FlaB binds TLR5 solely at the epitope, such that *Rh*FlaB PIM cannot bind full-length TLR5, we predicted that the FliC D0 would restore TLR5 binding to *Rh*FlaB PIM. As expected, swapping FliC D0 for the native *Rh*FlaB D0 rescues *Rh*FlaB PIM binding (Fig. 2C; lanes 6, 9). Taken together, these results show that FliC D0 allosterically binds TLR5, in direct contradiction to previous findings (*9, 11*). The additional binding site also explains why FliC binds full-length TLR5 more strongly than *Rh*FlaB (Fig. 2A).

The discovery of an allosteric TLR5 binding site in FliC prompted us to test its impact on TLR5 activation. We purified recombinant *Rh*FlaB chimera expressing the FliC D0 (*Rh*FlaB-FliC D0) and assayed TLR5 signaling. As expected from its greater ability to bind full-length TLR5, the chimeric flagellin is 100-fold more stimulatory than *Rh*FlaB with its native D0 (Fig. 2D). We hypothesized that the additional TLR5 binding site in the FliC D0 increases activity in part by enabling *Rh*FlaB-FliC D0 to interact with more TLR5 receptors than *Rh*FlaB.

TLR5 activation requires the formation of a symmetric 2:2 flagellin:TLR5 complex (*9, 22*). How this complex is assembled remains unclear, although it is widely stated that flagellin binding induces TLR5 dimerization (*11, 23*–*25*). Early cryo-EM work revealed, however, that human TLR5 forms asymmetric homodimers in the absence of flagellin, a conformation likely associated with multiple ligand binding sites, and thus a possible target of the FliC D0 domain (*26*). We investigated if TLR5 forms unliganded dimers by briefly treating TLR5-HA HEK cells with the membrane impermeable crosslinker BS^3^ (Fig. 2E). In addition to monomeric TLR5, we detected a higher molecular weight species consistent with the size of a TLR5 dimer. This result suggests that pre-formed TLR5 dimers are present on the cell surface, in the absence of ligand-induced dimerization that is commonly invoked for TLR5.

We tested the ability of FliC and *Rh*FlaB to interact with TLR5 dimers. We observed that FliC binds both monomer and dimer, while *Rh*FlaB only interacts with the monomer (Fig. 2E; lanes 3,4). The switching out of its native D0 for FliC D0 endows *Rh*FlaB the ability to bind the dimer (Fig. 2E; lane 5). This result suggests that the FliC D0 directly binds the ectodomain and activates TLR5 signaling in part by mediating binding to preformed TLR5 dimers. This observation supports our hypothesis that an allosteric binding site in its D0 enables FliC to interact with more receptors than *Rh*FlaB. Furthermore, the different oligomeric states of TLR5 targeted by FliC and *Rh*FlaB may partially account for the inability of *Rh*FlaB to antagonize FliC (Fig. 2F,G).

To assess how widespread silent flagellins are in the healthy human microbiome, we searched for the peptide sequences of silent flagellins using a published database comprising more than 33,000 flagellins (Methods). Candidate silent flagellins were selected based on their presence in human gut metagenomes and by similarity to the C-terminal region of *Rh*FlaB (Fig. S1, S3A). The list was further curated to exclude flagellins containing a basic residue (R/K) at position *Rh*FlaB aa478, based on our observation that *Rh*FlaB H478R shows a slight, but significant, increase in TLR5 stimulation (Fig. S3B,C). The final candidate silent flagellins are mostly, but not exclusively, from species belonging to the *Lachnospiraceae* family (75/78) (Fig. 1C,S4).

To verify whether these 78 candidate silent flagellins are indeed silent, we expressed them recombinantly to screen for both TLR5 signaling and TLR5^N14^ binding (Fig. 3A).

**Fig. 3.**
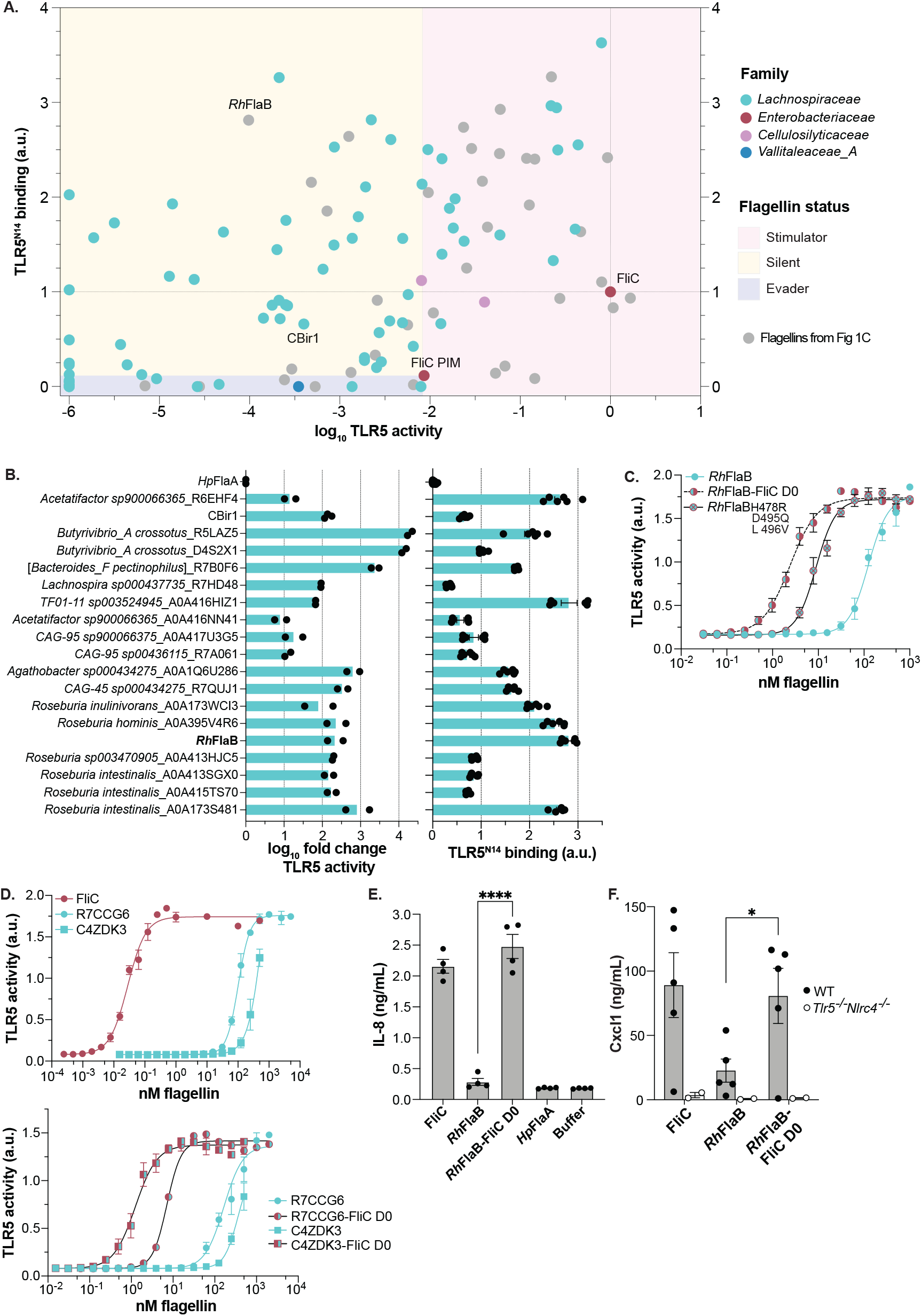
Silent flagellins are widespread among *Lachnospiraceae* that colonize the human gut. (**A**) Plotted are TLR5 activity vs TLR5^N14^ binding for flagellins with *Rh*FlaB-like C-terminal region: Each circle represents an individual flagellin and is colored by family-level taxonomy (GTDB); gray circles represent flagellins previously shown in Fig. 1C. TLR5^N14^ binding by AP-tagged candidates was performed as described in Fig.1; data shown represent mean for *n*≥3. TLR5 activity was calculated as described in Fig. 1C. Data represent mean from three independent experiments. All values are relative to FliC. See also Fig. S4. (**B**) Effect of FliC D0 on TLR5 activity: Native D0 domain was swapped for FliC D0 in a subset of *Lachnospiraceae* silent flagellins to generate chimeras, and TLR5 activity was measured as described in (A). Bar graph represents mean difference between wild-type and chimeric flagellin activity from at least two independent experiments. TLR5^N14^ binding of subset silent flagellins shown on right. (**C**) Effect of Flagellin cD0 residues on TLR5 activity: *Rh*FlaB cD0 was mutated at three sites to express residues present in FliC cD0, recombinantly purified, incubated with TLR5 HEK-Blue cells, and processed as described in Fig. 2B. (**D**) Activation of TLR5 by flagellins from human stool: Top two flagellins identified by proteomics were recombinantly purified and incubated with TLR5 HEK-Blue cells as described in Fig. 2B. See Tables S1, S2 and Data S1. (**E**) Flagellin-dependent responses in colonoids: Organoids derived from human colon were incubated with flagellins (10 nM) or buffer control for 18 hr. IL-8 levels in culture media were quantified by ELISA. Data shown represent mean ± SEM from one of two independent experiments. Significance between *Rh*FlaB and *Rh*FlaB-FliC D0 means was determined by unpaired, two-tailed t test (\*\*\*\**P*<0.0001). (**F**) TLR5-dependent responses in mice: Wild-type and *Tlr5*^*-/-*^*Nlrc4*^*-/-*^ mice were treated with indicated flagellins (10 ug) by intraperitoneal injection and Cxcl1 levels in blood were measured by ELISA. Significance between *Rh*FlaB and *Rh*FlaB-FliC D0 means was determined by unpaired, two-tailed t test (\**P*<0.05).

Compared to our initial screen (Fig. 1C), we were successful in enriching for silent flagellins: over half (44/78) are weaker TLR5 agonists and stronger TLR5^N14^-binders relative to FliC PIM (Fig. 3A, yellow region). Given its importance in Crohn’s disease, we additionally tested the flagellin CBir1, whose weak TLR5 agonism has been previously reported (*14, 27*). CBir1 binds TLR5^N14^, categorizing it as a silent flagellin. The remaining 34 candidates are equally distributed among stimulators (red region) and evaders (blue region).

To assess whether the mechanism is the same for *Rh*FlaB as for this new set of silent flagellins, we examined the impact of swapping in the FliC D0 domain on a subset representing a broad range of TLR5^N14^ binding strengths. While the magnitude differs among candidates, the FliC D0 universally increases TLR5 signaling for all silent flagellins tested, including CBir1 (Fig. 3B). Moreover, these silent flagellins belong to common taxa of the human gut microbiome, including multiple species of *Roseburia* (*17*). The FliC D0 does not affect TLR5 evasion by *Hp*FlaA, consistent with previous observations (*11*). To identify residues in the FliC cD0 responsible for increasing TLR5 activity, we looked for regions of low conservation between FliC and the subset silent flagellins. Substituting three amino acids in the *Rh*FlaB cD0 to the equivalent residues present in FliC increases TLR5 activity by *Rh*FlaB more than 10-fold (Fig. 3C, Table S1).

Given that flagellin is facultatively expressed, and that expression in the gut can vary depending on external factors (*13*), we assessed the presence of silent flagellins directly from healthy human stool. Endogenous flagellins were isolated using TLR5 as bait and identified by mass spectrometry. Peptides were searched against a custom flagellin database built from metagenome sequences generated from the same stool sample. Of the 12 flagellins identified, 10 are ascribed to *Lachnospiraceae* (Table S2, Data S1.). This is consistent with the taxonomic affiliation of the abundantly expressed flagellins in healthy humans (*18*) (Fig. S5A,B). We recombinantly expressed and purified the top two candidates to assay TLR5 signaling. Both flagellins weakly activate TLR5 with EC_50_ values greater than 100 nM (Fig. 3D; Table S1). But, similar to *Rh*FlaB, swapping in the FliC D0 for the native D0 profoundly increases their ability to stimulate TLR5.

We further verified that silent recognition occurs when TLR5 is endogenously expressed. We tested the effect of flagellins in 3D-cultured human colon organoids: FliC stimulates the secretion of IL-8, a pro-inflammatory cytokine produced downstream from TLR5 activation (Fig. 3E). IL-8 levels in *Rh*FlaB-treated organoids are similar to those of *Hp*FlaA- and buffer-treated controls, while organoids incubated with *Rh*FlaB-FliC D0 phenocopy FliC-treated organoids. Furthermore, mice injected with *Rh*FlaB have lower pro-inflammatory Cxcl1 cytokine levels compared to animals injected with FliC and *Rh*FlaB-FliC D0 (Fig. 3F). These results indicate that silent recognition of flagellin is not species-specific, and that the FliC D0 activates both human and mouse endogenously-expressed TLR5.

Our discovery of an allosteric activator of TLR5 in FliC suggests a mechanism by which this receptor can respond to minute levels of stimulatory flagellin. Commensal members of the gut microbiome, such as members of the *Lachnospiraceae*, can produce an array of flagellins that are silent, stimulatory, or evasive. *R. hominis* itself expresses several flagellins that fall into all three categories based on our binding and activation criteria (Fig. 4A; Tables S1,S3) (*19*). This within-species flagellin diversity reflects the flagellin diversity encoded broadly in human gut metagenomes, where all three types are detected (Fig. 4B; Fig. S6). We observed that non-Westernized metagenomes encode a greater proportion of all three flagellin types compared to Western metagenomes despite lower relative abundance of *Lachnospiraceae* (Fig. 4C; Fig. S5B-C). Of note, the decrease in flagellin abundance with Westernization is most pronounced for the silent flagellins (Fig. 4D).

**Fig. 4.**
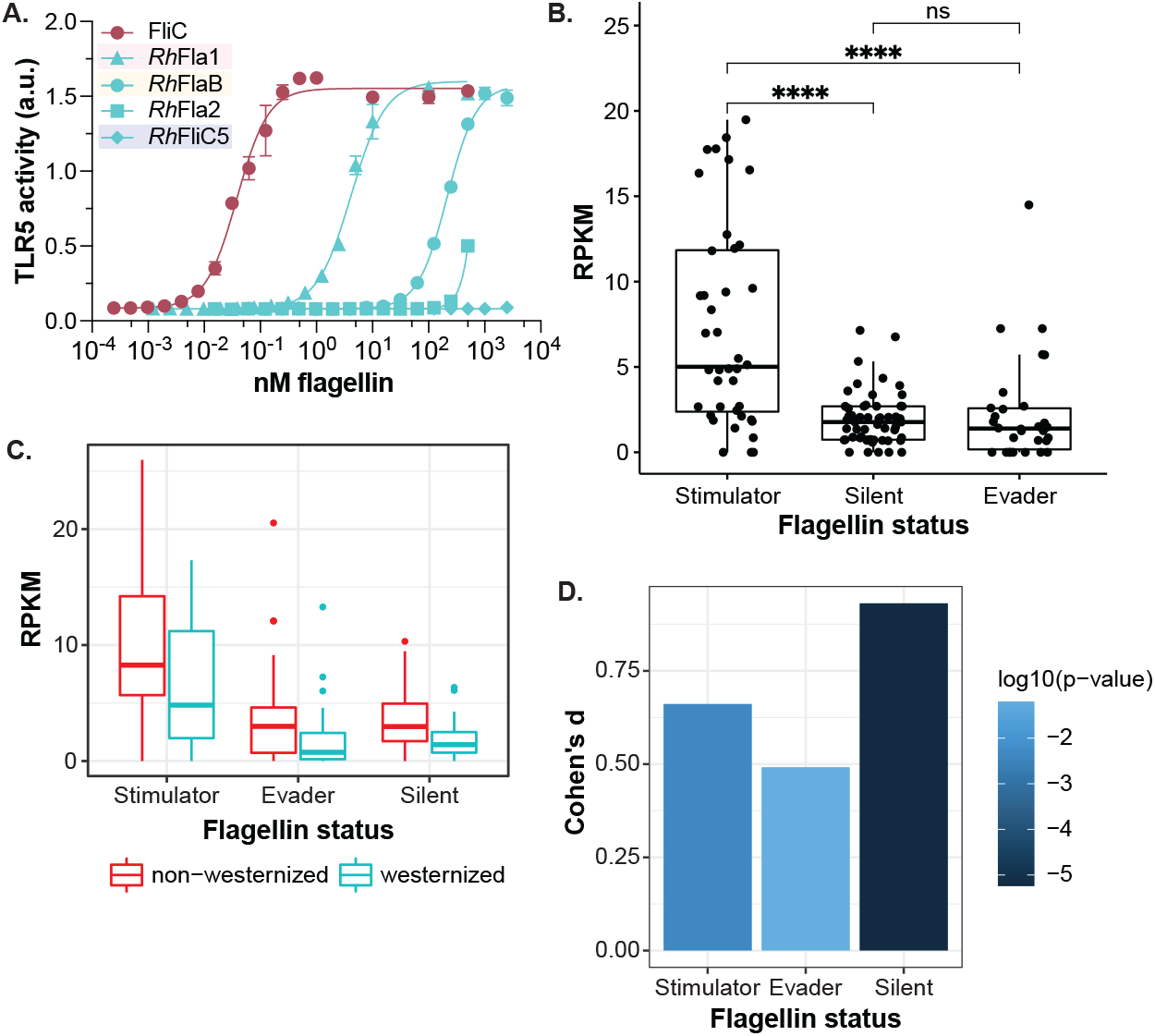
Silent flagellins are enriched in non-Westernized populations. (**A**) Within-species flagellin diversity: Flagellins encoded by *R. hominis* (Table S3) were recombinantly purified and incubated with TLR5 HEK-Blue cells as described in Fig. 2B. Flagellins are shaded to reflect status (Fig 3A, S2). See also Table S1. (**B**) Abundance of stimulator, silent, and evader flagellins: Boxplots show median reads per kilobase per million reads (RPKM) of flagellins in human metagenomes (*n*=1783). Significance determined by post hoc pairwise t-tests (ANOVA, type II, F = 25.3, P = 6.8e-10). (\*\*\*\**P*<0.0001; ns, not significant). See Fig. S6. (**C**) Median RPKM of flagellins by westernization status and flagellin status: See Fig. S6 and Data S2. (**D**) Effect sizes (Cohen’s D) and t-test P-values corresponding to westernization versus non-westernization comparisons within each flagellin status.

The understanding of how TLR5 interacts with its primary ligand, flagellin, has come mostly from the study of flagellins encoded by the *Pseudomonadota* (formerly *Proteobacteria)*, notably the pathogens *Salmonella* and *H. pylori*, and others such as *E. coli*. These studies led to the discovery of the TLR5 epitope, a conserved region on the flagellin nD1, whose binding is considered required for TLR5 recognition and subsequent stimulation. We show here that, in addition to the TLR5 epitope, the D0 of FliC allosterically binds TLR5, akin to a homotropic ligand. Our work indicates that in addition to these modes of interaction (*i*.*e*. recognition followed by activation versus non-recognition), a third mode, very common in commensal bacteria prevalent in the gut, allows bacteria to express flagellins that retain the TLR5 epitope without inducing a robust TLR5 response. While commensal bacteria also produce stimulatory and evasive flagellins (indeed all three types can be encoded in a single genome), our analysis of metagenomes indicates that silent flagellins are very common in the healthy human gut, and therefore represent a substantial, previously unappreciated, yet physiologically relevant population of TLR5 ligands with a novel mode of interaction.

Our current model proposes that TLR5 adopts different conformations, as evidenced by the presence of both monomeric and dimeric TLR5, and that flagellins have different affinities for these receptor states. While the FliC D0 confers high affinity for dimeric TLR5, silent flagellins have low affinity for this receptor state and are weak agonists relative to FliC as a result. However, this low affinity enables silent flagellins to activate TLR5 at high concentrations, in contrast to *Hp*FlaA. Our data further suggests that FliC binding to TLR5 complexes induces a conformational change, rather than receptor dimerization, similarly to what has been described for other TLRs (*28*).

Together, our work highlights how pattern recognition by TLR5 can occur without downstream signaling. By probing into the weak agonism of flagellins produced by commensal gut bacteria, we discovered a third class of flagellins, which contain the epitope recognized by TLR5 yet poorly activate the receptor. Allosteric activation of TLR5 allows the host to tolerate silent flagellins from commensal bacteria while remaining responsive to faint levels of stimulatory flagellin.

## Acknowledgments

Human colon organoids were from the Stanford Tissue Bank and kindly provided by Julia Y. Co from the laboratories of Manuel R. Amieva and Denise Monack. We thank Irina Droste-Borel and Boris Macek from the Proteome Center Tuebingen, Christa Lanz and Oliver Weichenrieder from the Genome Center at the MPI for Biology, and Ivan Hang and Jana Neuhold at the Vienna Biocenter Core Facilities (VBCF ProTech).

## Author contributions

Conceptualization: SJC, REL

Methodology: SJC, MB, AB, KP, ZMH, JCZ, NDY, REL

Investigation: SJC, MB, JCZ, AB, ZMH, KP, JZ, NDY

Visualization: SJC, NDY, AB Supervision: NDY, REL

Resources: DL, ZMH, YB, ATG, REL

Writing – original draft: SJC, JCZ, AB, NDY, DL, ATG, REL

## Data and materials availability

The raw sequence data of the sample used for proteomics analysis are available from the European Nucleotide Archive under study accession number PRJEB47632. Proteomics data are available from the Proteomics Identification Database (accession number pending).

## Supplementary Materials

Methods and Materials

Figs. S1 to S6

Tables S1 to S3

Data S1 to S2

References (*29*–*64*)

