## Supplementary Material for "Silent recognition of flagellins from human gut commensal bacteria by Toll-like receptor 5"

**This PDF file includes:**

Methods and Materials
Figs. S1 to S6
Tables S1 to S3
Captions for Data S1 and S2

**Other Supplementary Materials for this manuscript include the following:**

Data S1 and S2 [ProteinGroups, curMeta]

**Methods and Materials**

Selection of flagellins

The selection process of candidate flagellins is summarized in Fig. S1. Using the flagellin database from (*29*), we annotated database sequences with InterProScan (*30*) and only retained sequences having both Pfam domains PF00669 (flagellin N-terminal domain, 'ND') and PF00700 (flagellin C-terminal domain, 'CD'). Our filtered database comprised 33,051 flagellin protein sequences. To identify flagellins abundant in the human gut, the hits from metagenomes of healthy subjects from the inflammatory bowel disease multi-omics database of Lloyd-Price et al., 2019 (*18*) were filtered using the median of read counts as a cutoff. The taxonomy of those accessions was assigned using the taxonomizr R package (*31*) to obtain their taxids, and a further search of their taxids within the GTDB Database release 95 (https://data.ace.uq.edu.au/public/gtdb/data/releases/latest/). We searched the remaining accessions using Entrez Direct (*32*) in the NCBI Identical Protein Groups database (*33*) to obtain their assembly accession, which were further searched within the GTDB taxonomy. We ranked the resulting accessions by their read counts and randomly selected across the list to get a broad taxonomic representation of flagellins from the human gut, reducing the dataset to 44 candidates. We tested the random community assembly of the final candidates using different metrics: i) Cmean and Pagel's Lambda, implemented in the phylosignal R package (*34*) and the standardized effect size (SES) of ii) mean pairwise distance (MPD) and mean nearest taxon distance (MNTD) implemented in the picante R package (*35*). We retrieved the coding sequence (CDS) of each protein accession from the NCBI Identical Protein Groups database (*33*).

To assess the expression profile of flagellins in the human gut, we retrieved publicly available gut metatranscriptome data (*18*) (available at: https://ibdmdb.org/tunnel/public/HMP2/MTX/1750/products). We restricted our assessment to samples from the 26 healthy controls with associated metatransciptomes in (*18*), selecting one sample per subject. We filtered the downloaded HUMAnN2 tables (*36*) by retaining features annotated as flagellin, and whose abundance could be attributed to a specific taxon by HUMAnN2's tiered search. We further validated that the retained flagellins contained the N-terminal and C-terminal domains. HUMAnN2's tiered search provides NCBI taxonomy annotation; we re-annotated the contributing species by matching the NCBI taxonomy annotation to that of the Genome Taxonomy Database (GTDB) release 95 (*37*).

*Rh*FlaB-like candidates were identified using the flagellin database previously described. A truncated CD protein sequence of *Rh*FlaB (Accession: WP_014081191.1) was mapped against the database using DIAMOND blastp (*38*) with parameters `--max-target-seqs 0 --evalue $10^{-3}$ --very-sensitive`. The resulting 10,121 hits were filtered by i) the median of the length of the alignment (100aa), ii) the sequence identity (39%), iii) and the number of mismatches (n=60), which resulted in 979 protein sequences. We mapped the ND flagellin from *Salmonella* Typhimurium (Accession: AHA06007.1) against the selected hits with DIAMOND blastp using parameters `--max-target-seqs 0 --evalue 10-3 --very-sensitive`, which reduced the list to 919 protein sequences. The resulting list was manually curated by selecting those sequences that did not have either amino acids arginine or lysine in the position 478 of the alignment, leaving a total of 195 sequences.

We then sought to reduce the list of *Rh*FlaB-like flagellins to only those occurring in the human gut. We first mapped gut metagenomes from the inflammatory bowel disease multi-omics database (*18*) and the Franzosa et al., 2019 dataset (*39*) to our flagellin database using DIAMOND blastx with parameters `--max-target-seqs 1 --evalue 1e-3 --very-sensitive`. The resulting 5131 protein accessions were intersected with the 195 flagellins from the former step, resulting in 145 flagellins that meet the sequence composition criteria and are also present in the human gut. From these, we de-replicated at 99% sequence identity with CD-HIT (*40*, *41*) using the parameters `-c 0.99 -M 16000 -n 5`, which reduced the dataset to 85 protein accessions. We retrieved their CDSs from the NCBI Identical Protein Groups database (*33*). To avoid nucleotide sequence redundancy with the previously identified abundant flagellins, the lists of CDSs were merged, and the final nucleotide sequence list was de-replicated using CD-HIT with parameters `-c 0.99 -M 16000 -n 5.

Estimating abundance of silent, stimulator, and evader flagellins in human gut metagenomes

We used metagenomes from the curatedMetagenomicData dataset (*42*). Samples were selected based on the following criteria: i) shotgun metagenomes sequenced using the Illumina HiSeq platform with a median read length >95 bp; ii) available SRA accessions; iii) labeled as adults or seniors, or with a reported age ≥18 years; iv) without report of antibiotic consumption; v) without report of pregnancy; vi) non-lactating women; vii) labeled as 'healthy'. In cases of multiple samples per individual, one sample was randomly selected. The final dataset consisted of 1783 samples from 21 studies (see Data S2). Only the forward reads were used, and all metagenomes were subsampled to 1 million reads.

We mapped the metagenome reads to all stimulator, silent, and evader flagellins (n = 126) via DIAMOND blastx with the following parameters: --sensitive --iterate --evalue 1e-5 --query-cover 90. The results were used to calculate reads per kilobase of target per million reads (RPKM) for each flagellin in each sample. Cohen's D was calculated with the effsize R package (*43*).

Reagents and antibodies

Chemicals were purchased from Sigma, Carl Roth, or Thermofisher unless stated otherwise. Primary antibodies include mouse anti-Myc monoclonal 4A6 (Cat. No. 05-724) and rat anti-HA monoclonal 3F10 (Roche Cat. No. 11867423001). Secondary antibodies were from LI-COR (IRDye 800CW Goat anti-Rat IgG and IRDye 680RD/800CW Goat anti-Mouse IgG).

$TLR5^{N14}$ binding to AP-tagged flagellins

Unless stated otherwise, flagellin (prey) and $TLR5^{N14}$ (bait) constructs were cloned into a modified pLIB vector backbone containing an N-terminal BiP insect signal peptide (MKLCILLAVVAFVGLSLG) and a C-terminal Strep-II affinity tag as described previously (*44*). Flagellin constructs were further modified to include a C-terminal alkaline phosphatase (AP) tag. The $TLR5^{N14}$ bait construct was generated by inserting the first 14 LRRs of human TLR5 flanked by an N-terminal cap and C-terminal variable leucine repeat adaptor sequence followed by human IgG-Fc and a 6XHis tag. Constructs were codon-optimized for expression in *Drosophila melanogaster* and assembled using Gibson Assembly® Master Mix (NEB E2611). Plasmids were transformed into NEB 5- Competent E. coli (Cat. No. C2987), cultured in 5 mL LB supplemented with carbenicillin (100 ug/mL), and purified using the ZymoPURE Plasmid Miniprep kit (Zymo Research Cat. No. D4211).

Sequence verified constructs were transformed into DH10EMBaCY cells to generate recombinant baculovirus genomes (bacmids) as described previously (*44*, *45*). Bacmids were transfected into SF9 insect cells using the FuGENE® 6 Transfection Reagent (Promega Cat. No. E2691), and expanded to generate an initial (V0) viral stock. Two successive rounds of viral amplification were performed through 1:100 v/v inoculations of 25 mL SF9 cell cultures with baculovirus-containing supernatant, each lasting 72 hr. The supernatant from the second culture (V2) was used to induce protein production. Protein production was optimized for each construct through expression trials in Hi5 cells using inoculation ratios from 1:10 to

1:1000. Proteins were harvested after 48 hr except for FliC PIM and *Hp*FlaA, which were harvested after 72 hr.

Secreted prey proteins in supernatant were collected following centrifugation (800 rcf, 5 min) and filter purified. Bait protein was obtained from Hi5 cell pellets by ultrasonication in lysis buffer (50mM HEPES, 300mM NaCl, 5% Glycerol, 0.1% Tween-20, 5mM 2-Mercaptoethanol, 1x Protease-inhibitor (Serva, 39107)), then filter purified. Protein expression was confirmed by immunoblotting with Strep-Tactin® conjugated to horseradish peroxidase (HRP) (IBA, 2-1502-001).

Flagellin candidates from human commensals were PCR amplified from pETM11 bacterial expression vectors using Phusion High-Fidelity DNA Polymerase (Thermo Scientific, Cat. No. F530S). The insert was cloned into the pECIA14 vector using Gibson assembly (NEBuilder® HiFi DNA Assembly Master Mix, Cat. No. E2621). Insect cell protein expression was performed as previously described (*46*, *47*) with minor modifications. AP-tagged flagellins were expressed via transient transfection in *Drosophila melanogaster* Schneider 2 (S2) cells using Expres$^2$ TR Transfection Reagent (Expres$^2$ion Biotechnologies). During transfection, the cells were shifted from 27°C to 25°C. Protein expression was induced with 1mM $CuSo_4$ 24 hr post-transfection. Supernatant media were collected 72 hr post-induction. Protease inhibitors (cOmplete, EDTA free, Roche, Cat. No. 5056489001) and 0.02% $NaN_3$ were added to the media. The expressed proteins were confirmed by immunoblotting using anti-Flag-HRP (Sigma Aldrich, Cat. No. A8592) antibody.

Each quantification and plate-assay contained *Rh*FlaB*,* FliC*,* and *Hp*FlaA as controls. AP-tagged flagellins were quantified by incubating 25 µL of each prey sample in 100 µL BluePhos® Microwell Substrate (Seracare Cat. No. 5120-0059) for 30 min at room temperature. Prey concentrations were normalized to *Rh*FlaB based on their relative response to the BluePhos® Substrate by dilution in PBS-T (PBS + 0.1% v/v Tween-20). $TLR5^{N14}$-Flagellin binding was assayed following a previously described method (*46*), with some modifications. Flagellin proteins and $TLR5^{N14}$ were pre-diluted 1:10 in PBS-T, then mixed by rocking for 2 hr at 4ºC. 96-well Pierce™ Protein A Coated plates (ThermoFisher Cat. No. 15132) were activated by two washes with 200uL PBS-T. 200 uL of the $TLR5^{N14}$-Flagellin mixtures were added to each well, and incubated overnight at 4ºC. Each sample had 3 replicates per plate, alongside corresponding flagellin-only wells as controls. AP activity was quantified using BluePhos® Microwell Substrate, with the absorbance measured at 650 nm after 3 hr. Relative $TLR5^{N14}$-Flagellin binding strength was quantified by subtracting the background absorbance from flagellin-only wells from mixed wells, then normalized to FliC. Negative values were set to zero.

<u>Preparation of Myc-tagged flagellins</u>

cDNAs encoding flagellins were commercially synthesized (IDT) with an N-terminal Myc tag and cloned into the pETM11 vector downstream from a 6x-histidine tag (NEBuilder HIFi DNA assembly) prior to transformation in TOP10 competent cells (Invitrogen). Mutated flagellins and flagellins expressing the D0 domain from *Salmonella* FliC (aa1-41 and aa456-494) were generated using Gibson cloning. Plasmid inserts were confirmed by Sanger sequencing and transformed into ClearColi BL21(DE3) cells (Lucigen).

Unless stated otherwise, ClearColi-transformed cells were grown at 37°C in LB supplemented with kanamycin (50 ug/mL) to $OD_{600}$ 0.4-0.6/mL. Protein expression was induced following the addition of 0.1 M IPTG and cells were incubated for 2.5 hr at 37°C. Cells were harvested by centrifugation (3400 x g, 15 min) and pellets were resuspended in 1:100 volume lysis buffer (10 mM Tris·HCl pH 8, 8M urea, 100 mM $NaH_2PO_4$). Cells expressing *H. pylori* FlaA (or FlaA-FliC D0) were grown at 25°C for 4 hr following induction then lysed in 10 mM Tris·HCl pH 8, 6 M guanidine HCl, 100 mM $Na_2HPO_4$. Cells were lysed for 2 hr at room temperature and the soluble fraction was collected after centrifugation (16000 x g, 20 min). These lysates were used to screen for TLR5 activity in Figs. 1C and 3A,B.

For flagellin purification, cleared lysates were incubated with Ni-NTA agarose (1:4 volume; Qiagen) for 2 hr at room temperature. Agarose matrix was then washed once with two column volumes (CV) lysis buffer then washed twice with two CVs lysis buffer pH 6.3 and twice with two CVs lysis buffer pH 5.9. Ni-NTA-bound proteins were eluted from the matrix with 0.5 CV lysis buffer pH 4.5. Eluates were dialyzed at 4°C in 20 mM Tris·HCl pH 8, 300 mM NaCl, 5 mM $MgCl_2$ with the exception of *H. pylori* FlaA, CAZDK3, CAZDK3-FliC D0, and R7CCG6-FliC D0, which were dialyzed in 50 mM HEPES pH 7.4, 114 mM NaCl, 1.5 mM $Na_2HPO_4$. Purified proteins were quantified by BCA (Pierce) and aliquots stored at -80°C.

For mouse experiments, flagellins lacking a Myc-tag were purified as described above then passed through a Polymyxin B agarose column (Sigma) prior to BCA quantification.

TLR5 HEK Blue activity assay

HEK-Blue hTLR5 cells (InvivoGen) were grown in 5% $CO_2$ at 37°C in medium (GlutaMAX DMEM, 10% FBS (Gibco)) containing selection antibiotics (100 ug/mL Zeocin; 30 ug/mL blasticidin) to 90% confluence. Cells were detached using pre-warmed PBS pH 7.4 (Gibco), resuspended in medium ($1 \times 10^5$ per mL), and distributed in 96-well plates (180 uL/well). Purified flagellins were serially diluted in PBS and 20 uL added in triplicate to HEK cells for a final volume of 200 uL/well. Plates were incubated for 18 hr (5% $CO_2$, 37°C) then 20 uL medium from each well was added to 180 uL QUANTI-Blue solution (InvivoGen). Absorbance at 635 nm was measured after 30 min, with the exception of Fig. 3C, which was measured after 80 min. $EC_{50}$ values for purified flagellins were calculated by plotting absorbance values against flagellin concentrations in Prism 9 (GraphPad) and performing weighted, non-linear regression analysis. Unless stated otherwise, no constraints were set. $LogEC_{50}$ values were averaged.

Flagellin candidates were screened for TLR5 activity (Figs. 1C, 3A) as described above with the following modifications. HEK cells were plated 190 uL/well and bacterial lysates expressing flagellins were serially diluted in PBS such that the final volume in the assay ranged from 1 to $1 \times 10^{-6}$ uL. Lysates were added in duplicate to HEK cells to a final volume of 200 uL/well. The relative amount of flagellin expression was determined by quantifying the Myc signal in 0.1 uL lysate (diluted 10-fold in PBS) by immunoblotting. TLR5 activity was determined by plotting absorbance values against bacterial lysate volume in Prism 9 and calculating $EC_{50}$ values from weighted, non-linear regression analysis with the following constraints: Hill slope set to 1 and top less than or equal to a shared value. Activity was normalized to protein expression by calculating $EC_{50}$ multiplied by Myc signal relative to FliC. Values were negative $\log_{10}$ transformed. Candidates with activity less than empty vector control were assigned $\log_{10}$ TLR5 activity values equal to -6.

Full-length TLR5-flagellin pull-down assays

hTLR5-HA HEK cells (InvivoGen) were incubated in 5% $CO_2$ at 37°C in medium (GlutaMAX DMEM, 10% FBS (Gibco)) supplemented with the selection antibiotic blasticidin (10 ug/mL). Cells were grown to 90% confluency in T75 flasks (Grenier), detached with pre-warmed PBS pH 7.4, and harvested by centrifugation (3400 x g, 3 min). Pellets were resuspended in 1 mL pre-chilled lysis buffer (25 mM Tris·HCl pH 7.5, 1% v/v Igepal CA-630, 100 mM NaCl, 5 mM imidazole, 10% v/v glycerol) supplemented with 1x HALT protease inhibitors (Thermo Scientific).

Cells were incubated on ice for 15 min followed by centrifugation at 4°C (855 x g, 20 min). Supernatant was collected and protein levels were quantified by BCA. For each reaction, 500 ug cell lysate was diluted in 300 uL lysis buffer and incubated with 150 pmol Myc-tagged flagellin or buffer control for 2 hr at 4°C. TALON resin (Takara) was washed twice with PBS pH 7 and twice with lysis buffer prior to resuspension in PBS; 20 uL washed resin was added to each reaction. Following 2 hr incubation at 4°C, resin was pelleted at 4°C (855 x g, 1 min) and washed three times with 750 uL lysis buffer supplemented with protease inhibitors (cOmplete, mini, EDTA-free). Bound fraction was eluted from resin with 20 uL lysis buffer containing 200 mM imidazole and transferred to fresh tubes. Samples (25 ug input and eluted fractions) were analyzed by gel electrophoresis followed by immunoblotting with antibodies against Myc and HA.

For pull-downs with crosslinked lysates, hTLR5-HA HEK cells were plated in Nunc EasYdishes (Thermofisher Cat. No 150460) to 70% confluence. Immediately prior to $BS^3$ treatment, cells were washed with 1 mL PBS then incubated with 2.33 mM $BS^3$ crosslinker (Pierce Cat No. 21580) in 1.5 mL PBS for 30 sec on ice. Reaction was quenched with the addition of 1 M Tris·HCl pH 8 to final concentration 50 mM. Crosslinker was removed and cells were scraped into pre-chilled lysis buffer (750 uL) then processed as described above with the following modification: input sample was incubated with anti-HA resin (Roche Cat. No. 11815016001) for 2 hr at 4°C and bound proteins were eluted with 20 uL HA peptide (1 mg/mL in TBS; Thermofisher Cat. No. 26184) for 15 min at 37°C.

### Immunoblotting

Protein samples were diluted in 1X Laemmli loading buffer (50 mM Tris·HCl pH 6.8, 2% w/v SDS, 2% v/v glycerol, 0.05% w/v Bromophenol Blue, 2.5% v/v BME) and incubated at 95°C for 5 min. Samples and Chameleon Duo Pre-stained protein ladder (LI-COR) were loaded in 4-12% NuPAGE Bis-Tris protein gels (Invitrogen) and ran in a Mini Gel Tank (Invitrogen) with MES buffer (50 mM MES, 50 mM Tris pH 7.3, 0.1% w/v SDS, 1 mM EDTA) at 125 V. Following electrophoresis, gels were transferred to nitrocellulose membranes (Pierce) at 10V for 70 min at 4°C in transfer buffer (25 mM Bicine, 25 mM Bis-Tris pH 7.2, 1 mM EDTA, 10% v/v methanol) using the Mini Blot Module (Invitrogen). Membranes were blocked for 30 min with 5% w/v milk in TBS-T (19 mM Tris pH 7.6, 137 mM NaCl, 2.7 mM KCl, 0.1% v/v Tween-20) then incubated in primary antibody for 1 hr at room temp or overnight at 4°C. Membranes were washed 3X for 5 min with TBS-T then incubated with secondary antibody for 30 min at room temp. Membranes were washed 3X prior to imaging on the Odyssey CLx (LI-COR). Primary and secondary antibodies were diluted in TBS-T with 5% w/v milk to 0.1 ug/mL (anti-HA), 0.5 ug/mL (anti-Myc), and 0.05 ug/mL (IRDyes).

Band intensities from immunoblots were quantified in Image Studio (LI-COR). For pull-downs, the fraction of TLR5 bound to flagellin was calculated by first subtracting the background HA signal then dividing by the Myc signal and normalizing to TLR5 monomer bound to FliC.

### Stool sample preparation for proteomics

A stool sample was previously obtained from a healthy adult female (Cornell University Institutional Review Board, protocol number 1106002281). Stool (166 mg) was resuspended in PBS pH 7.4 supplemented with protease inhibitors to 0.03% w/v and vortexed at max speed for 20 min. Lysates were cleared following centrifugation at 13,700 x g for 10 min at 4°C. Protein levels in the supernatant were quantified by BCA, aliquoted, and stored at -80°C. TLR5-HA HEK lysates were prepared from ~6.5 x $10^6$ cells as described above in lysis buffer lacking imidazole and incubated with 90 uL anti-HA resin for 2 hr at 4°C. Beads were washed

3x with lysis buffer prior to incubation with 1.8 mg stool proteins for 2 hr at 4°C. Prior to incubation, stool proteins were boiled for 5 min at 98°C and diluted in lysis buffer to final volume 300 uL. After incubation, beads were washed 3x with lysis buffer and bound proteins were analyzed by mass spectrometry.

Flagellin database for proteomics

We extracted total DNA from the above stool sample using the Qiagen PowerSoil kit following the manufacturer's instructions. Library preparation and sequencing were performed as described above. After quality control, the sequencing depth of the sample was 210,366,585 paired reads. We mapped the sequencing reads to our curated flagellin database using DIAMOND v.2.0.9 (*48*) using parameters `--evalue 1e-3 --ultra-sensitive`. The abundance of mapped reads was transformed to reads per kilobase (RPK). We assigned taxonomy to the flagellin hits by matching the NCBI protein ID and its associated assembly to GTDB r95. The final database contained 1028 flagellin sequences.

NanoLC-MS/MS analysis and MS data processing

Proteins were eluted from the washed beads and purified with a 12% NUPAGE Novex Bis-Tris Gel (Invitrogen). Tryptic in-gel digestion of proteins was performed as described previously (*49*) and extracted peptides were desalted using C18 StageTips (*50*). Eluted peptides were subjected to LCMS/MS analysis.

Peptide analysis was performed on an Easy-nLC 1200 system coupled to an Exploris 480 mass spectrometer (Thermo Fisher Scientific) as described elsewhere (*51*) with slight modifications: peptides were injected onto the column in HPLC solvent A (0.1% formic acid) at a flow rate of 500 nl/min and subsequently eluted with a 227 min segmented gradient of 10–33-50-90% HPLC solvent B (80% ACN in 0.1% formic acid). During peptide elution the flow rate was kept constant at 200 nl/min.

In each scan cycle, the 20 most intense precursor ions were sequentially fragmented using higher energy collisional dissociation (HCD) fragmentation. For both precursors and fragment ions the maximal injection time mode was set to auto and the AGC target to standard. Precursor masses with charge states between 2 and 6 were selected with a resolution of 60,000 at minimum intensity of 400,000 at a scan range of 300 to 1750m/z. They were excluded from further selection for 30 s. HCD collision energy for peptide fragmentation was set to 28%, resolution for MS2 scans was 30,000.

MS data were processed with MaxQuant software suite version 1.6.7.0 (*52*) and database search was performed using the Andromeda search engine (*53*), a module of the MaxQuant. MS/MS spectra were searched against a *Homo sapiens* database obtained from Uniprot (released 07.10.2020, 97,795 entries), a database containing 1028 flagellin sequences from various organisms (European Nucleotide Archive study accession number PRJEB47632), and a database consisting of 246 commonly observed contaminants. In database search, full tryptic specificity was required and up to two missed cleavages were allowed. Carbamidomethylation of cysteine was set as fixed modification, whereas oxidation of methionine and acetylation of protein N-terminus were set as variable modifications Mass tolerance for precursor ions was set to 4.5 ppm and for fragment ions to 20 pm. Peptide, protein and modification site identifications were reported at a false discovery rate (FDR) of 0.01, estimated by the target/decoy approach (*54*). For protein group quantitation a minimum of two quantified peptides were required. All search parameters were kept to default values except for the following. Minimal peptide length of 5 amino acid was required, and the iBAQ algorithm was used to estimate quantitative values by dividing the sum of peptide intensities of all detected peptides by the number of theoretically observable peptides of the matched protein (*55*). See Data S1.

Organoid experiments

Organoids derived from human colon were cultured in Basement Membrane Extract (Cultrex PathClear Reduced Growth Factor, Type 2) and growth medium (Advanced Dulbecco's modified Eagle medium/F12 supplemented with 50% v/v L-WRN conditioned medium described in (*56*), 1 mM HEPES, 1x glutamax, 1x B27, 1 mM N-acetylcysteine, 10 nM gastrin, 50 ng/mL EGF, 10 mM nicotinamide, 500 nM A83-01, 10 µM SB202190,10 µM Y27632, 250 nM CHIR99021) for 4 days in 96-well plates. Organoids were incubated for 18 hr in the presence of flagellins (10 nM) or buffer control diluted in growth medium at 37°C. IL-8 cytokine levels were quantified from 20 uL medium according to manufacturer's instructions (R&D Systems Human IL8 DuoSet ELISA).

Mouse experiments

All animal studies were performed at Georgia State University under an approved animal protocol (IACUC # A17047). Eight-week-old female C57BL/6 WT and $Tlr5^{-/-}$/$Nlrc4^{-/-}$ mice, bred at Georgia State University, were treated with 10 µg FliC, *Rh*FlaB, or *Rh*FlaB-FliC D0 by intraperitoneal injection. Two hr later, blood was collected from these mice via retrobulbar intraorbital capillary plexus, and serum was isolated by centrifugation at 4 °C using MiniCollect serum separator tubes (Greiner Bio-One). Cxcl1/Kc in serum were quantitated by Duoset ELISA kits from R&D Systems according to the manufacturer instructions.

General data analysis and visualization

Data was processed, analyzed, and visualized with snakemake (*57*), conda (*58*), and R (*59*) and the following R packages: dplyr (*60*), tidyr (*61*), ggplot2 (*62*), and ggpubr (*63*). PDB structure was color-coded using PyMOL (*64*).

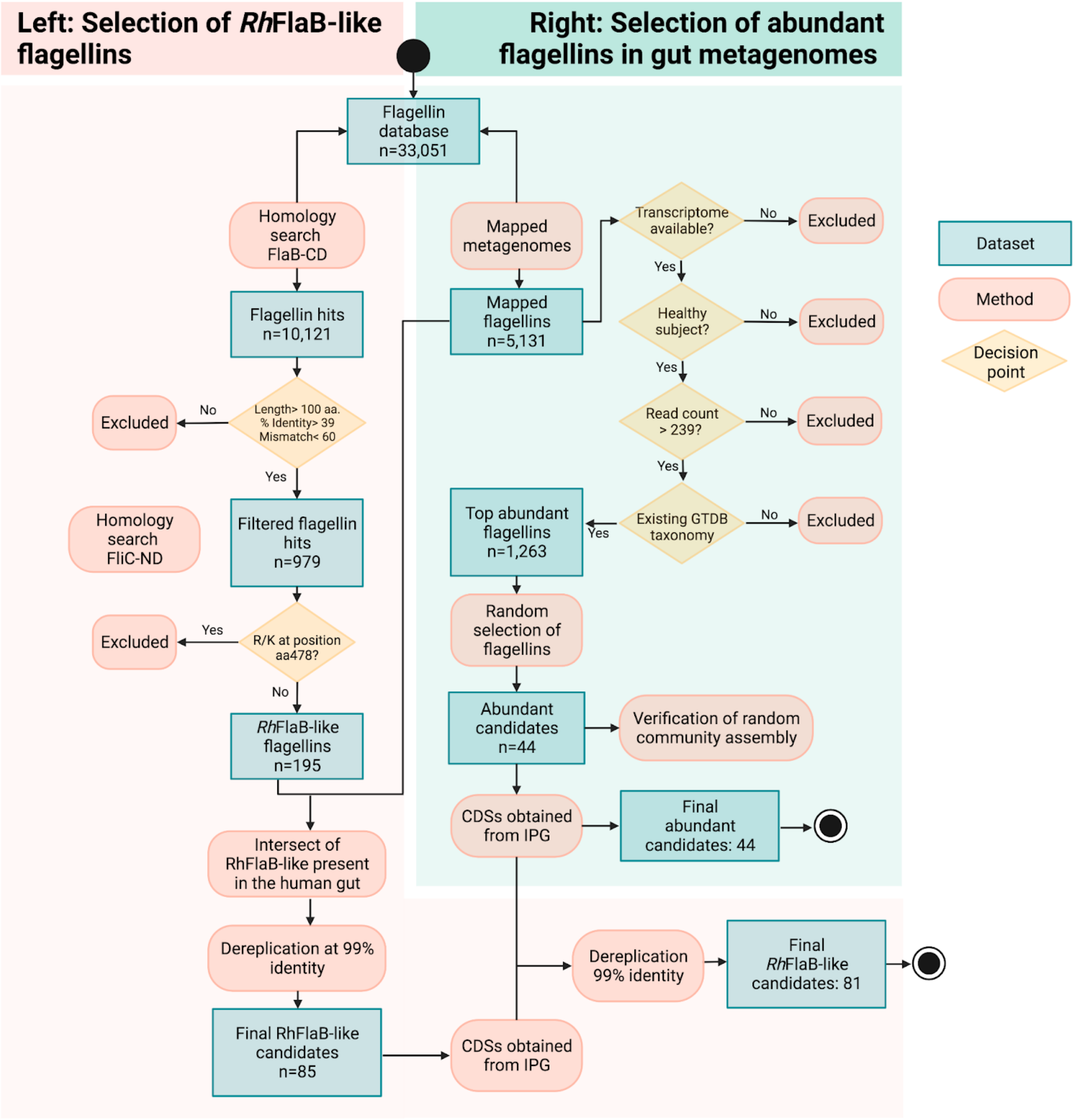


**Fig S1. Selection of flagellin candidates.** The flowchart illustrates the strategy used to identify abundant flagellins ($n$ = 44) and *Rh*FlaB-like flagellins occurring in the human gut ($n$ = 81). **Left:** *Rh*FlaB-like flagellins were identified by homology search of truncated C-terminal domain (CD) and N-terminal domain (ND) of FlaB of *Roseburia hominis* and FliC of *Salmonella Typhimurium*, respectively. Only *Rh*FlaB-like flagellins occurring in human gut metagenomes were kept. **Right:** Abundant flagellins from gut metagenomes of healthy subjects were randomly selected among flagellins with a read count over the median and an existing taxonomy on GTDB v95. The coding sequences (CDSs) of *Rh*FlaB-like candidates were de-replicated with the CDSs of previously identified abundant flagellins to avoid redundancy. Six candidates were excluded due to poor expression in insect cells resulting in 41 highly abundant and 78 *Rh*FlaB-like candidates. See Methods for additional details.

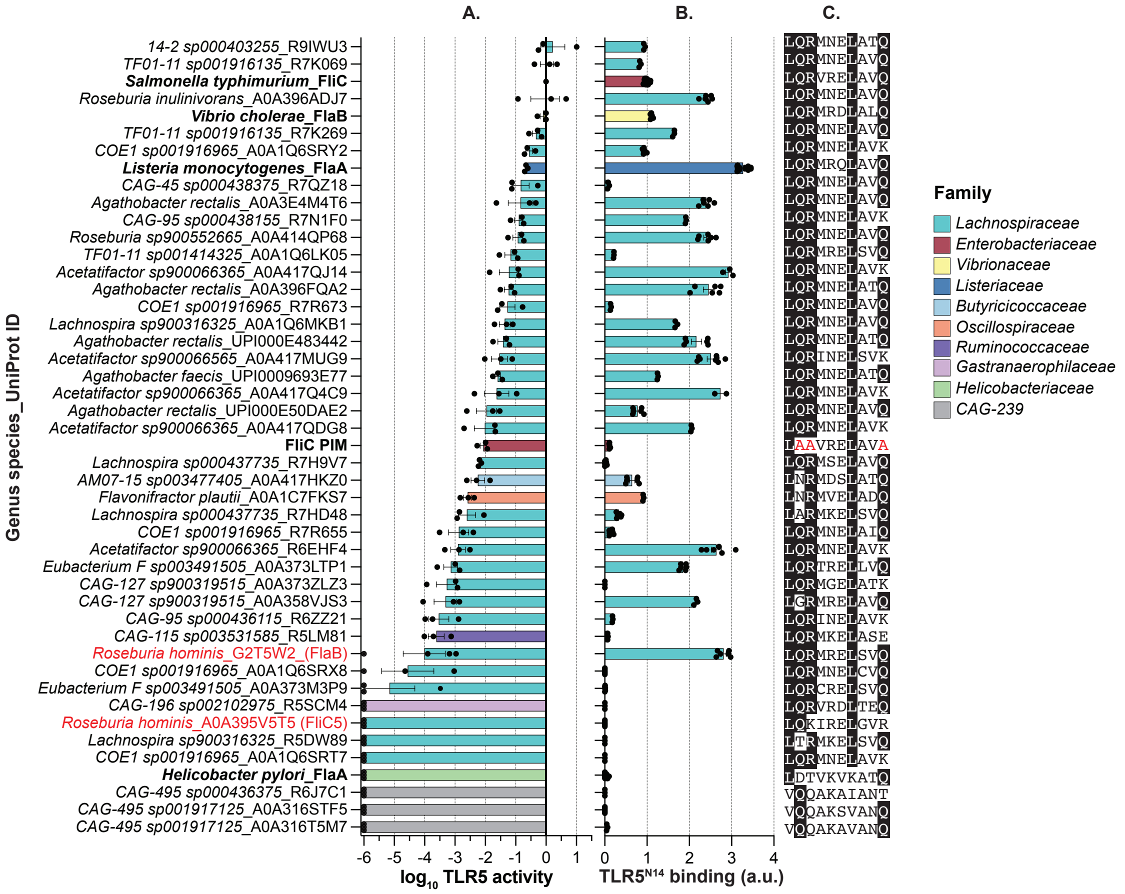


**Fig. S2**. **TLR5 recognition and activation by abundant flagellins from human gut.** (**A**) Activation of TLR5 by commensal flagellins from initial screen in Fig 1C. Myc-tagged flagellins ($n$=46) were expressed in bacterial lysates and incubated with TLR5 HEK-Blue cells. TLR5 activity represents negative $EC_{50}$ normalized to Myc expression. Data are mean ± SEM from three independent experiments. All values are normalized to FliC. (**B**) Flagellin binding to $TLR5^{N14}$. AP-tagged flagellins were incubated with $TLR5^{N14}$ bait and AP activity was quantified. Data are mean ± SEM for $n \geq 3$ and normalized to FliC. (**C**) TLR5 epitope sequences. Conserved residues required for TLR5 recognition are shaded black; residues mutated in FliC PIM are colored red. Flagellins produced by *R. hominis* (*Rh*FlaB and *Rh*FliC5) are red; flagellins from pathogens are shown in bold. Bar color indicates family-level taxonomy (GTDB). See also Fig. 1C.

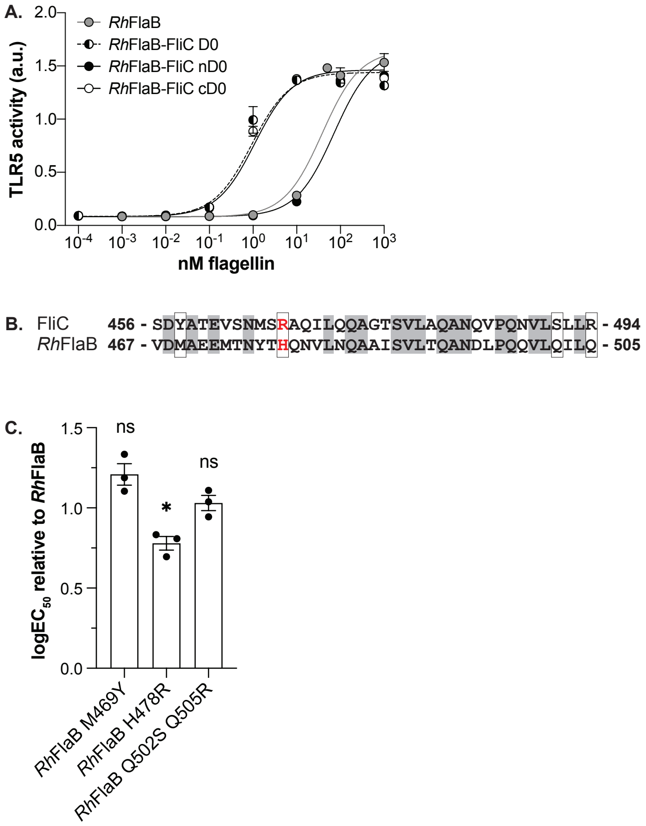


**Fig. S3**. **FliC C-terminal D0 (cD0) domain is necessary and sufficient for increased activation of TLR5.** (**A**) Effect of FliC nD0 and cD0 domains on *Rh*FlaB-dependent TLR5 activity. *Rh*FlaB chimeras were generated by substituting the native nD0 and cD0 with those of FliC. Chimeras were recombinantly purified and incubated with TLR5 HEK-Blue cells to measure activity. Data are mean ± SEM for $n$=3 and represent one of two independent experiments. Curve-fitting by weighted, non-linear regression analysis with Hill slope constrained to 1. (**B**) Non-conserved residues in flagellin cD0. Alignment of FliC and *Rh*FlaB cD0 domain amino acid sequences. Shaded areas denote identical residues and boxes indicate mutants tested in (C). *Rh*FlaB H478 and FliC R467 are colored red. (**C**) Effect on TLR5 activity when *Rh*FlaB is mutated to express cD0 residues present in FliC. *Rh*FlaB mutants were recombinantly purified and incubated with TLR5 HEK-Blue cells. Data shown are mean $logEC_{50}$ ± SEM ($n$=3) of mutants normalized to wild-type *Rh*FlaB as determined by weighted, non-linear regression analysis with Hill slope constrained to 1 . Significance was calculated by one sample t test (*$P$<0.05; ns, not significant).

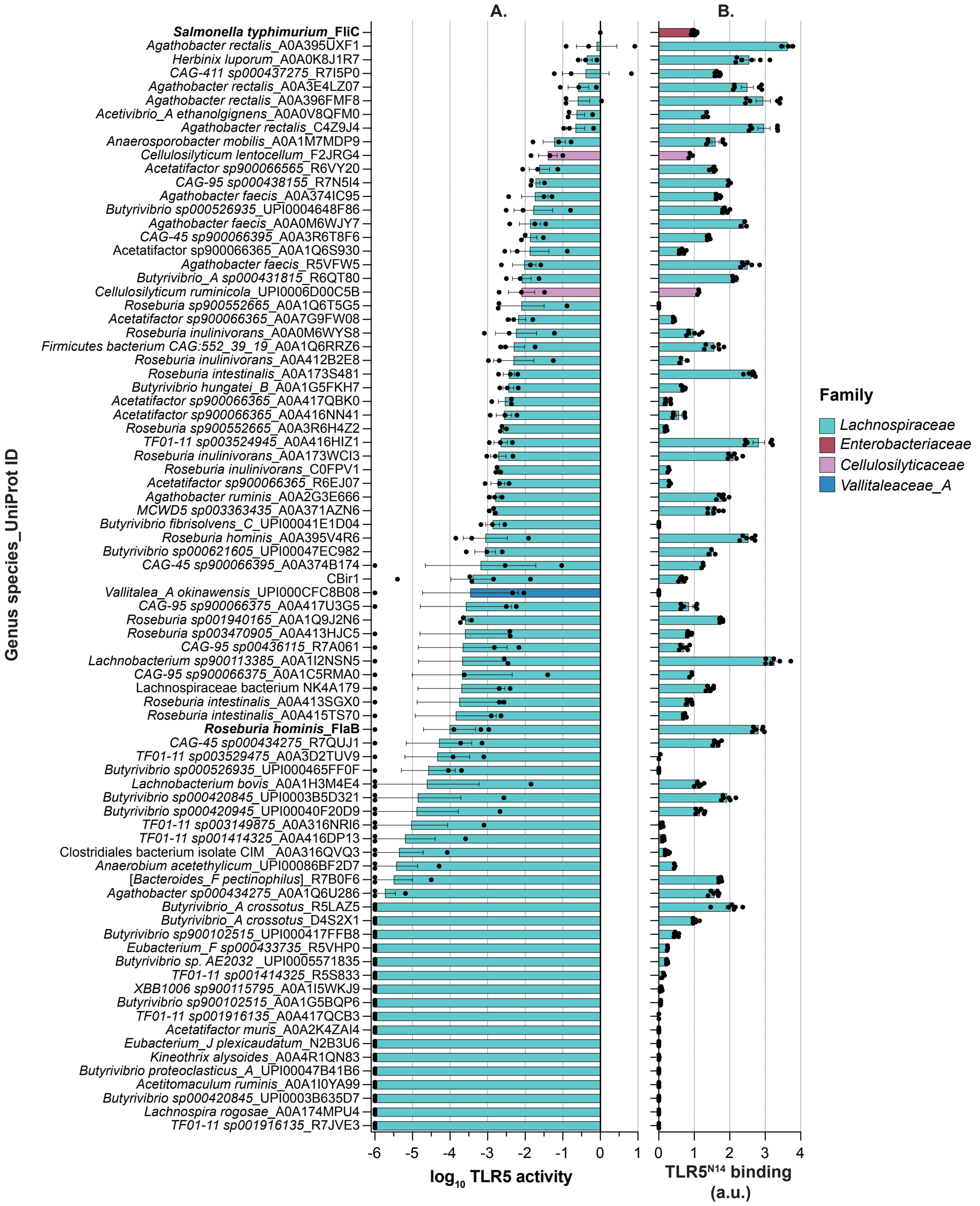


**Fig. S4. TLR5 recognition and activation by *Rh*FlaB-like flagellins**. (**A**) Activation of TLR5 by *Rh*FlaB-like flagellins from Fig 3A. TLR5 activity was measured as described in Fig S2A. Data are mean ± SEM from three independent experiments. All values are normalized to FliC. (**B**) Flagellin binding to TLR5$^{N14}$. AP-tagged flagellins were incubated with TLR5$^{N14}$ bait and AP activity was quantified. Data are mean ± SEM for $n \geq 3$ and normalized to FliC. Bar color indicates family-level taxonomy (GTDB). FliC and *Rh*FlaB are shown in bold. See Fig. 3A.

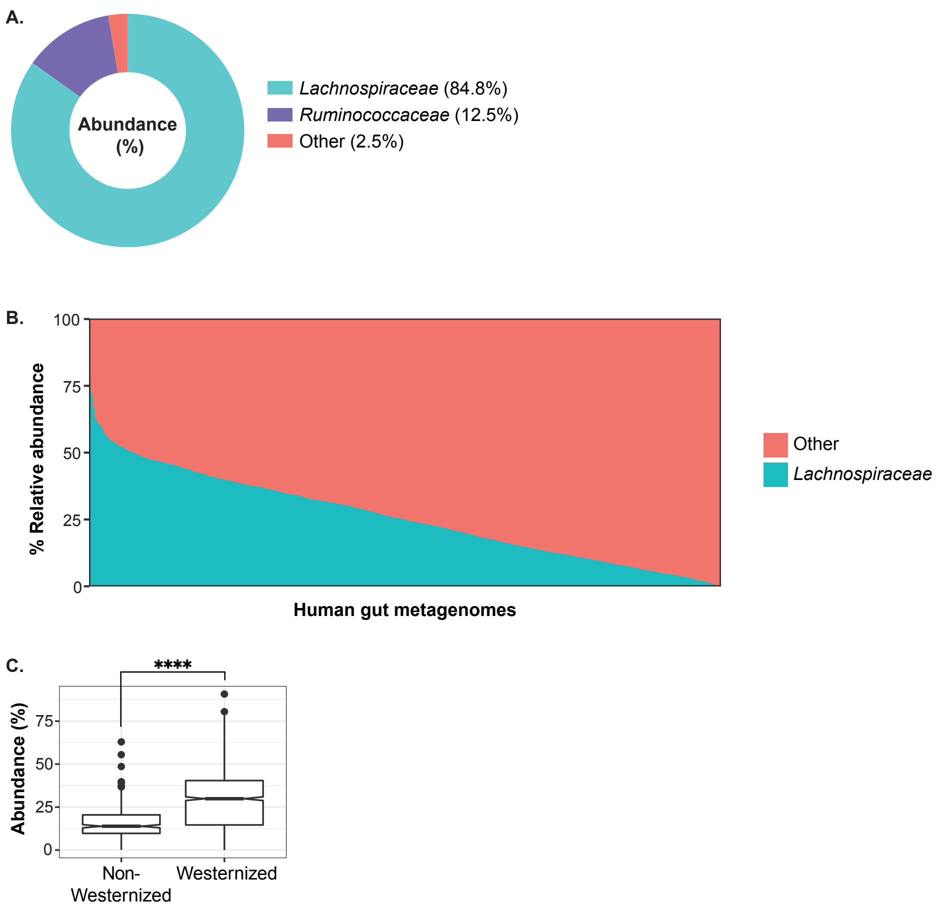


**Fig. S5**. ***Lachnospiraceae* are dominant producers of flagellin in the human gut.** (**A**) Relative abundance of flagellin gene transcripts from metatranscriptomes of 207 healthy controls from ref *18*, as shown by family-level taxonomy (GTDB). (**B**) Relative abundance of *Lachnospiraceae* in human gut metagenomes from the curatedMetagenomeData collection (*n* = 1783). Only samples labeled as "healthy" were included. The dataset comprised 21 studies and individuals from 18 countries. Metagenome profiling was conducted via Kraken2 and Bracken with a custom reference database generated from the Genome Taxonomy Database (GTDB) Release 95 (*35*). (**C**) Median relative abundance of *Lachnospiraceae* in Westernized vs non-Westernized populations from (B). Significance between medians determined by Wilcoxon (*****P*<2.2e-16).

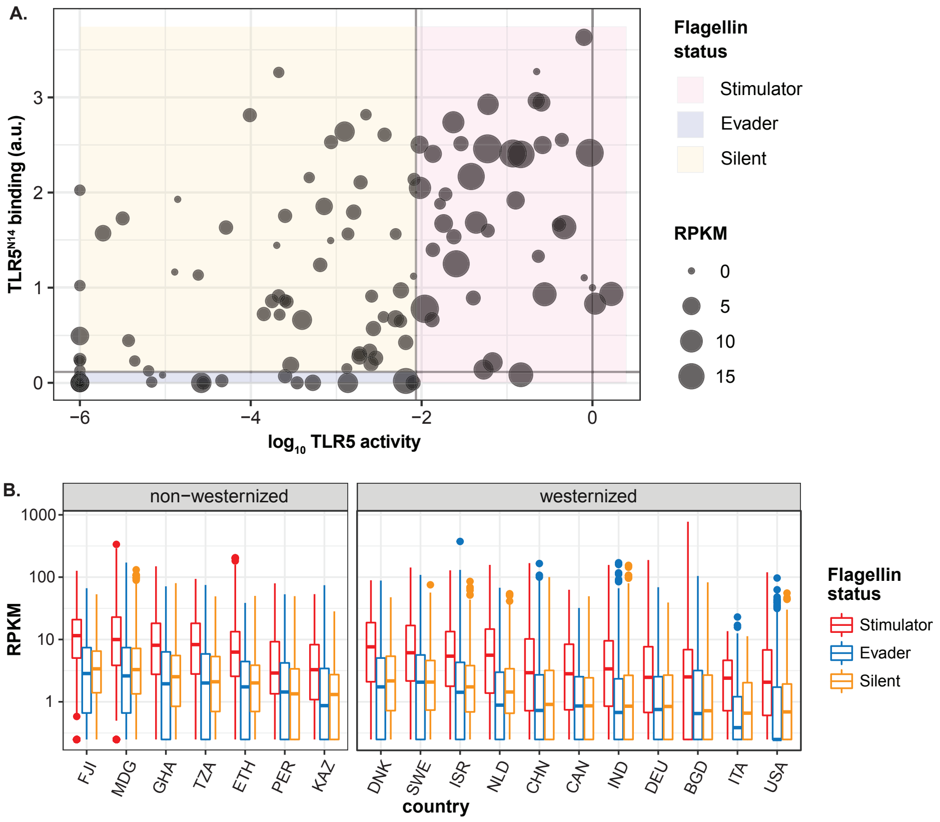


**Fig. S6. Active flagellins are more abundant than Evader and Silent flagellins among healthy human populations.** (**A**) The activity, binding, and median reads per kilobase per million reads (RPKM) for each flagellin. RPKM values were calculated by mapping metagenome Illumina reads from "healthy" individuals in the curatedMetagenomics dataset (*n* = 1783). See Data S2. (**B**). Median RPKM for each flagellin, grouped by country (ISO 3166-1 codes), westernization status, and flagellin status.

| **Taxonomy** | **Flagellin** | **$EC_{50}$ (nM)** | **log$EC_{50}$ ± SEM** |
|---|---|---|---|
| *Salmonella typhimurium* | FliC | 0.0644 | -1.19 ± 0.0867 |
| | FliC PIM (Q89A R90A Q97A) | 9.44 | 0.975 ± 0.0936 |
| *Roseburia hominis* | FlaB | 138.8 | 2.14 ± 0.0564 |
| | FlaB-FliC D0 | 2.53 | 0.403 ± 0.0632 |
| | FlaB H478R D495Q L496V | 9.66 | 0.985 ± 0.0335 |
| | Fla1 | 3.37 | 0.528 ± 0.0926 |
| | Fla2 | >1000 | 3.25 ± 0.460 |
| | FliC5 | >1000 | 3.31 ± 0.199 |
| *Agathobacter rectalis* | C4ZDK3 | 487.2 | 2.69 ± 0.0613 |
| | C4ZDK3-FliC D0 | 1.01 | 0.004 ± 0.0915 |
| *Eubacterium_F sp003491505* | R7CCG6 | 114.2 | 2.06 ± 0.0535 |
| | R7CCG6-FliC D0 | 3.95 | 0.596 ± 0.257 |

**Table S1.** $EC_{50}$ values for purified recombinant flagellins.

| Taxonomy | Flagellin (UniProt ID) | % coverage | Maxquant score | TLR5 epitope |
|---|---|---|---|---|
| *Agathobacter rectalis | WP_012744006.1 (C4ZDK3) | 40.4 | 238 | LQRMNELATQ |
| *Eubacterium_F sp003491505 | CDD75417.1 (R7CCG6) | 29.2 | 131 | LQRARELVVQ |
| *Roseburia intestinalis | CDA56948.1 (R6AYN5) | 25.2 | 68.0 | LQRMNELATQ |
| *Roseburia inulinivorans | WP_055168195.1 (A0A173SD30) | 23.9 | 60.0 | LQRMNELAVQ |
| *UBA9502 sp003480315 | SCG91213.1 (A0A1C5KT57) | 8.4 | 12.4 | LQRMNELAVK |
| *Lachnospira eligens_A | RHL68652.1 (A0A174Z6V7) | 15.1 | 9.66 | LQRMNELATQ |
| *14-2 sp000403845 | EOT27140.1 (S0JEY7) | 9.2 | 9.23 | LQRMNELATQ |
| *Roseburia intestinalis | RHA58738.1 (A0A3R6A0K5) | 25.9 | 8.56 | LQRMNELATQ |
| unclassified Clostridium | WP_117780849.1 (UPI000E553912) | 8.5 | 6.99 | LQRMGELATK |
| *TF01-11 sp001916135 | CDE69670.1 (R7K069) | 8.5 | 6.70 | LQRMNELAVQ |
| *Roseburia inulinivorans | RHA89587.1 (A0A413TXC5) | 25.9 | 5.91 | LQRMNELATQ |
| Clostridium hydrogeniformans | WP_027633903.1 (UPI000485A193) | 5.9 | 5.87 | LQRMRELSVQ |

**Table S2.** Endogenous flagellins identified directly from human stool by mass spectrometry using TLR5 as bait. Asterisk denotes members of the *Lachnospiraceae* family. Conserved residues in the TLR5 epitope required for recognition are underlined.

| **Gene** | **Old gene name** | **Protein** | **TLR5 epitope** |
|---|---|---|---|
| RHOM_RS00690 | RHOM_00665 | Fla1 | LQRMNELATQ |
| RHOM_RS15365 | RHOM_15820 | FlaB | LQRMRELSVQ |
| RHOM_RS00845 | RHOM_00820 | Fla2 | LDRMVELTTQ |
| RHOM_RS00990 | RHOM_00975 | FliC5 | LQKIRELGVR |

**Table S3.** Flagellins encoded by *R. hominis.* Conserved residues in the TLR5 epitope required for recognition are underlined.

**Data S1. ProteinGroups (separate file)**

All protein groups identified in the processed raw proteomics files (spectra). Each row contains the group of proteins that could be assigned to a set of identified peptides that were processed with a 1% false discovery rate.

**Data S2. curMeta (separate file)**

All relevant metadata associated with the curatedMetagenomics dataset samples used in this work. The country identifiers are ISO 3166-1. All data was obtained from the full curatedMetagenomics dataset.
